# On the inconsistent treatment of gene-protein-reaction rules in context-specific metabolic models

**DOI:** 10.1101/593277

**Authors:** Miguel Ponce-de-León, Iñigo Apaolaza, Alfonso Valencia, Francisco J. Planes

## Abstract

With the publication of high-quality genome-scale metabolic models for several organisms, the Systems Biology community has developed a number of algorithms for their analysis making use of ever growing –omics data. In particular, the reconstruction of the first genome-scale human metabolic model, Recon1, promoted the development of Context-Specific Model (CS-Model) reconstruction methods. This family of algorithms aims to identify the set of metabolic reactions that are active in a cell in a given condition using omics data, such as gene expression levels. Different CS-Model reconstruction algorithms have their own strengths and weaknesses depending on the problem under study and omics data available. However, after careful inspection, we found that all of these algorithms share common issues in the way GPR rules and gene expression data are treated. The first issue is related with how gapfilling reactions are managed after the reconstruction is conducted. The second issue concerns the molecular context, which is used to build the CS-model but neglected for posterior analyses. To evaluate the effect of these issues, we reconstructed ∼400 CS-Models of cancer cell lines and conducted gene essentiality analysis, using CRISPR–Cas9 essentiality data for validation purposes. Altogether, our results illustrate the importance of correcting the errors introduced during the GPR translation in many of the published metabolic reconstructions.

---

With the publication of high-quality genome-scale metabolic models (GSMs) for several organisms, the Systems Biology community has developed a plethora of algorithms for their analysis making use of ever-growing *omics* data (Heirendt *et al*., 2019). In particular, in human metabolism, the reconstruction of the first genome-scale model RECON1 (Duarte *et al*., 2007) promoted the development of Context-Specific Model (CS-Model) reconstruction methods (Opdam *et al*., 2017). This family of algorithms aims to identify the catalogue of metabolic reactions involved in a cell in a given condition using *omics* data, commonly gene expression levels. CS-Models are particularly suitable for studying metabolic differences between human tissues and are widely used in the area of human health (Uhlen *et al*., 2016), with applications including the prediction of metabolic vulnerabilities in cancer (Agren *et al*., 2014) and the inference of biomarkers in Alzheimer and diabetes (Geng and Nielsen, 2017), among others. Currently, we have dozens of these CS-Models available in different public repositories, usually stored under the Systems Biology Markup Language standard (Hucka *et al*., 2003).

An essential component of CS-Model reconstruction algorithms are Gene-Protein-Reaction (GPR) rules, which include the information about how genes relate to protein complexes and isozymes as well as the reactions they catalyze. GPR rules are expressed in the form of logical equations and allow us to define active/inactive reactions by mapping expression data onto them. This set of reaction states is used by reconstruction algorithms to extract the final CS-Model, which involves the reactions mapped as active plus a small set of inactive reactions automatically added to fill gaps in the network (gap-filling reactions). The relevance of GPR rules for CS-Model reconstruction algorithms can be more clearly observed in the Supplementary Information, where a general workflow is depicted.

Different CS-Model reconstruction algorithms have their own strengths and weaknesses depending on the problem under study and *omics* data available. In this direction, Opdam and collaborators performed an extensive benchmark of CS-Model algorithms and found that none particular method outperforms the others (Opdam *et al*., 2017). However, after careful inspection, we found that all of these algorithms share a common “bug” in the way GPR rules and gene expression data are treated when reconstructing CS-Models.

**The first issue we encountered is related to how gap-filling reactions are managed in the reconstruction process.** Model extraction algorithms may add reactions classified as inactive to fill gaps in the CS-Model. Figure 1A represents a toy model with four reactions: two categorized as active, {1, 4}, and two as inactive, {2, 3}. In this example, the CS-model includes the active reactions and the reaction 2 for gap filling. Importantly, the decision to include the gap-filling reaction 2 implies to update our assumption about the state of genes involved in such reaction, which in this case means to update the state of gene B to *active*. In addition, if the state of gene B is updated, this change must be propagated through the CS-model. Following the example in Figure 1A, propagating the change on the state of gene B implies that reaction 3 becomes active and thus it should be included in the consolidated CS-Model. We found that this consolidation step is not performed by the published CS-Model reconstruction algorithms, and that **neglecting the GPR consolidation step will worsen the model predictions by overestimating the effect of gene knockouts.**

**Figure 1.**
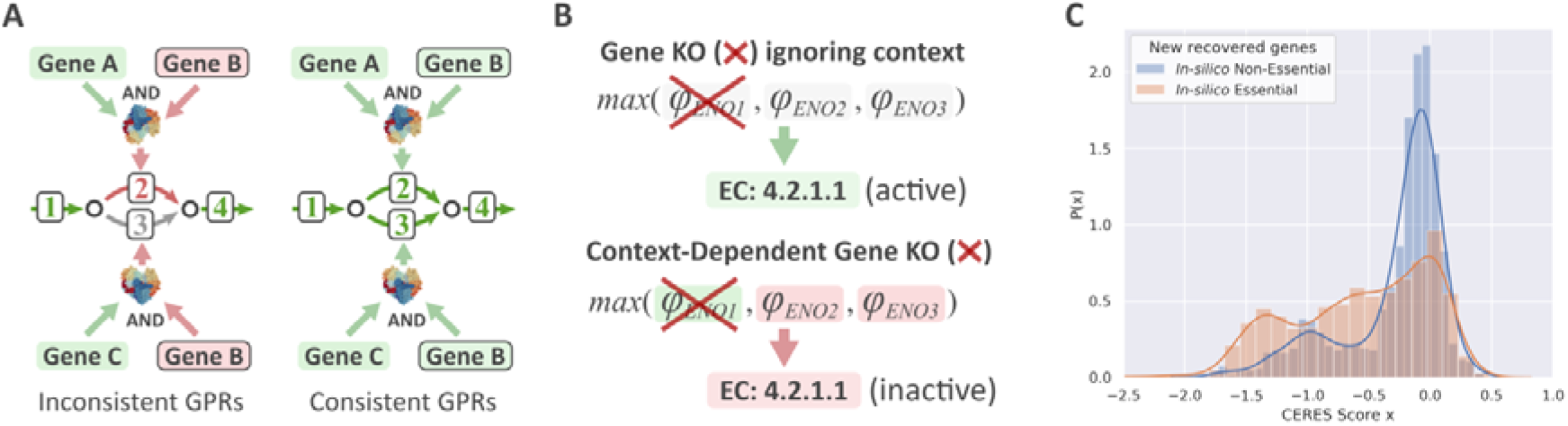
Identified errors in context-specific metabolic model reconstruction. **(A)** Illustration of the consequences of not managing the gap-filling reactions properly. **(B)** Illustration of the consequences of not taking the molecular context into account. **(C)** The distribution of CERES essentiality scores (Meyers *et al*., 2017) for recovered essential (orange) and non-essential (blue) genes. The recovered essential genes are those genes which are predicted non-essential by standard GIMME (Becker and Palsson, 2008) but essential when the errors in (A) and (B) are amended. The recovered non-essential genes are those genes which are predicted essential by standard GIMME (Becker and Palsson, 2008) but non-essential when the errors in (A) and (B) are amended. Red and green coloring refers to inactive and active genes, respectively. The green arrows correspond to active reactions, the red arrow refers to an inactive reaction which has been selected as gap-filling and the gray arrow corresponds to an inactive reaction which has not been selected as gap-filling.

The second and more striking issue is that **the molecular context used to reconstruct the CS-model is usually not included as part of the CS-model.** If the expression confidences used to infer the reaction states (*active/inactive)* are left aside after the reaction mapping, part of the information used to reconstruct the CS-Model is lost and, therefore, the model formulation will be incomplete. In the example of Figure 1B, the *enolase* reaction was set as active based on its expression scores and GPR rule. However, if the GPR of *enolase* is evaluated without the expression scores, it is not possible to guess which genes are supporting the *enolase* activity. Moreover, without the information about gene expression, the default hypothesis will be that all the genes included in the CS-Model are *active*. Thus, analyzing a CS-Model without considering the context can affect the predictions (see Figure 1B). **Specifically, we found that this inconsistent treatment of the CS-Model leads to down-estimating the effect of gene knockouts.**

In order to evaluate the effect of this flaw, we reconstructed ∼400 CS-Models for cell lines from the Cancer Cell Line Encyclopedia (Barretina *et al*., 2012) and conducted a gene essentiality analysis. The CS-Models were reconstructed using Gene Expression Barcode to classify genes as *active*/*inactive* (McCall *et al*., 2014), Recon3D as the reference model (Brunk *et al*., 2018) and GIMME as the extraction algorithm (Becker and Palsson, 2008). We amended the output of GIMME to account for the issues discussed above, using a simple and effective approach, which is fully detailed in the Supplementary Information. **Summarizing the results over all the cell lines, our approach found 3160 essential genes not predicted by standard GIMME and discarded 1061 essential genes predicted by standard GIMME** (Supplementary Data). These results clearly indicate that ignoring the molecular context has drastic effects on the *in-silico* predictions.

To validate these results, for each cell line we gathered CRISPR–Cas9 essentiality data from DepMap (Tsherniak *et al*., 2017) corrected using CERES essentiality score (Meyers *et al*., 2017). Figure 1C shows the CERES scores’ distributions for the aforementioned 3160 non-essential genes becoming essential (orange curve) and 1061 essential genes becoming non-essential when GIMME considered context (blue curve). **As expected, the first group is significantly enriched in DepMap essential genes** (one tailed Mann-Whitney test p-value= 9.96·10^-41^), demonstrating the practical importance of the correct treatment of GPR rules and molecular context.

Altogether, our results illustrate the importance of the errors introduced during the GPR translation in many of the published metabolic reconstructions. To overcome this issue, we advocate for a strict control of the specific molecular context during the translation of the GPR rules to CSMs. To that end, the **existing CSM reconstruction algorithms and storage standards should be modified to be GPR consistent and provide the molecular context.** Here, we showed the positive results in the performance of GIMME when this limitation was corrected and similar results are expected with other algorithms.

