## Supplementary Text for "On the inconsistent treatment of gene-protein-reaction rules in context-specific metabolic models"

### Building a Context-Specific Model (CS-Model)

Figure S1 shows the process of building a Context-Specific Model (CS-Model). First, the expression levels of the genes involved in metabolic processes are converted into a expression score or confidence, usually by discretizing the expression levels into two (or more) states, *e.g.* *active*/*inactive*. In order to classify the enzymatic activities of the reference model, the expression scores are mapped onto the reactions following the Gene-Protein-Reaction (GPR) rules. As a result, each reaction of the reference model has an assigned state (*active/inactive*) (Fig. S1A). Finally, a reconstruction algorithm (*e.g.* GIMME, Fig. S1B) takes a reference model and the *active* reactions list from the previous step and extracts the Context-Specific Model (CS-Model) (Fig S1C).

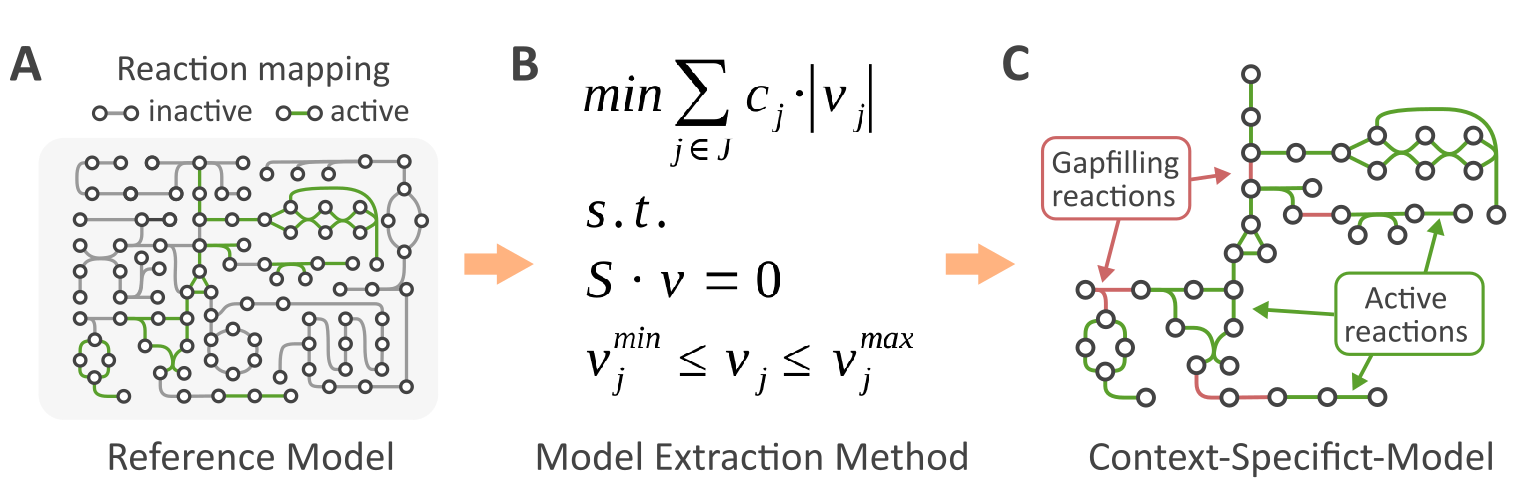

***Figure S1. Context-Specific Reconstruction.*** ***(A)*** *Mapping gene expression levels into reactions in the reference metabolic model.* ***(B)*** *A model extraction method is applied to the reference network integrating the expression levels.* ***(C)*** *Example CS-Model.*

A key input data for CS-model reconstruction algorithms (discussed in the main text) are GPR rules, which are a set of Boolean equations that relate genes and reactions (Fig. S2). They include *AND*/*OR* operators that represent enzyme-complexes/isozymes, respectively. Thanks to them, we can move from gene to reaction scores. In particular, in order to obtain the confidence score of a particular reaction from the scores of its associated genes, for a reaction catalyzed by different isozymes (*OR* operator), the maximum confidence score of the associated genes is considered (Fig. S2A). On the other hand, in the case of enzyme complexes (*AND* operator), the minimum confidence score of its associated genes is considered (Fig. S2B). More complex cases are solved by applying the same approach recursively (Fig. S2C).

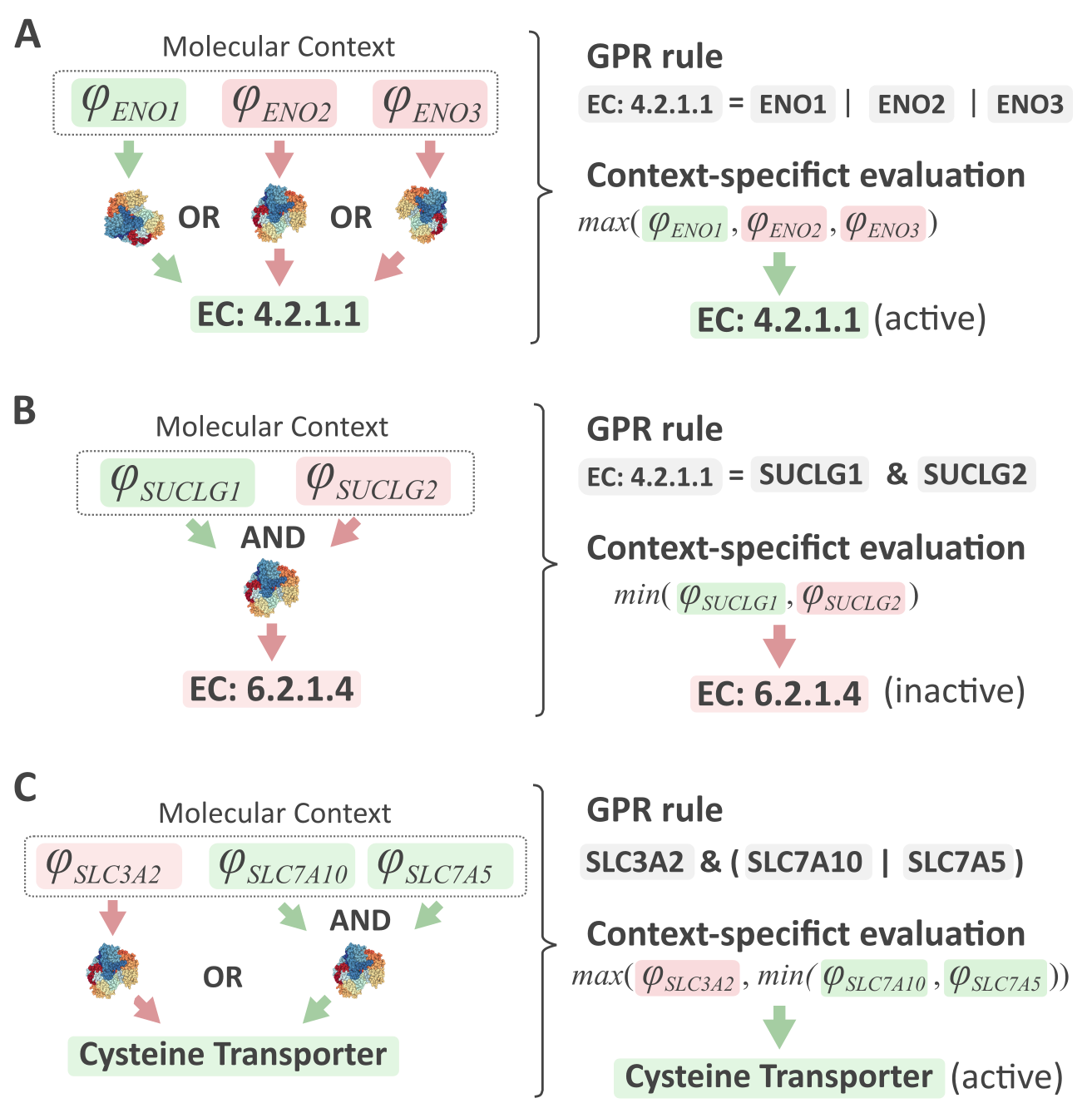

***Figure S2. Mapping expression levels onto reactions.*** ***(A)*** *Metabolic reaction with a simple OR GPR rule (isozymes).* ***(B)*** *Metabolic reaction with a simple AND GPR rule (enzymatic complex).* ***(C)*** *Metabolic reaction with a complex AND/OR GPR rule.*

### Corrections to GIMME and gene essentiality analysis to correctly consider GPR rules and molecular context

In the main manuscript we explain that approximately 400 different cell lines from the Cancer Cell Line Encyclopedia (Barretina *et al*, 2012) have been included in this study (Supplementary Table 1). Exactly the same analysis has been performed to each of the cell lines as described below.

This work involves two main steps: (1) the contextualization of the reference metabolic network Recon3D_3.01 (Brunk *et al*, 2018) for each cell line included in the study by integrating the transcriptomics data (Supplementary Table 1) and (2) the performance of the gene essentiality analysis.

#### Context-specific reconstruction using GIMME

First, we downloaded the gene expression data from the Gene Expression Omnibus portal (Edgar *et al*, 2002) and processed it following the Gene Expression Barcode pipeline (McCall *et al*, 2014). With the normalized gene expression levels we applied GIMME (Becker & Palsson, 2008) to extract a context-specific subnetwork using Recon3D_3.01 (Brunk *et al*, 2018) as the reference metabolic network. The formulation of the optimization problem underlying the GIMME method is shown below:

$$min\sum_{j\in J} c_{j}\cdot\left| v_{j} \right|$$

$$s.t.:$$

$$S\cdot v=0$$

$v_{j}^{min}\leq v_{j}\leq v_{j}^{max}$

$v_{BIO}\geq v_{BIO}^{max}$

where:

$$c_{j}=\left\{ \begin{matrix} T-e_{j} if T>e_{j} \\ 0 if T\leq e_{j} \end{matrix} \right\}$$

being $T$ the expression level above which genes/reactions are treated as expressed; and $e_{j}$ the expression value for reaction $j$. On the other hand, $S$ stands for the stoichiometry matrix; $v$ is the flux vector; $v_{BIO}$ is the flux through the biomass reaction; $v_{BIO}^{max}$ is the maximum flux through the biomass reaction obtained after solving FBA. Note that the gene expression levels must be transformed to the reaction level by means of the Gene-Protein-Reaction (GPR) rules.

By solving the linear program shown above, we obtained a list of reactions that are included in the CSM. After adding all the expressed reactions and those whose expression level is unknown to the aforementioned list of reactions, we obtained a reconstructed network for each cell line. Note that these “conventional reconstructions” are calculated at the reaction level and that this fact introduces certain errors in the analyses due to the complexity of the GPR rules or the missing information, as explained in the main manuscript. Therefore, in order to avoid the concerns raised, we fixed these “conventional reconstructions” as follows.

Genes whose expression level is above $T$ are considered active in the reconstruction. For each of the low confidence reactions included in the “conventional reconstruction” (gapfilling reactions), we took their associated genes and then iteratively decreased $T$ until the reaction becomes active. In other words, we calculated the minimum number of genes per reaction, in descending order of the expression level, to be added in the reconstruction for the reaction under study to be active. Note that all those genes whose expression level is unknown are given the maximum value. At this point, we obtained a list of genes which were included in the reconstruction and another set of genes which were not.

The next step consisted in translating the “gene-level” reconstruction to the reaction level by means of the GPR rules. Note that all the reactions included in the “conventional reconstructions” were included in these “corrected reconstructions”. However, some new ones arise as a consequence of complex GPR rules and the addition of genes that are necessary to activate the low confidence reactions involved in the reconstruction (Fig. S3).

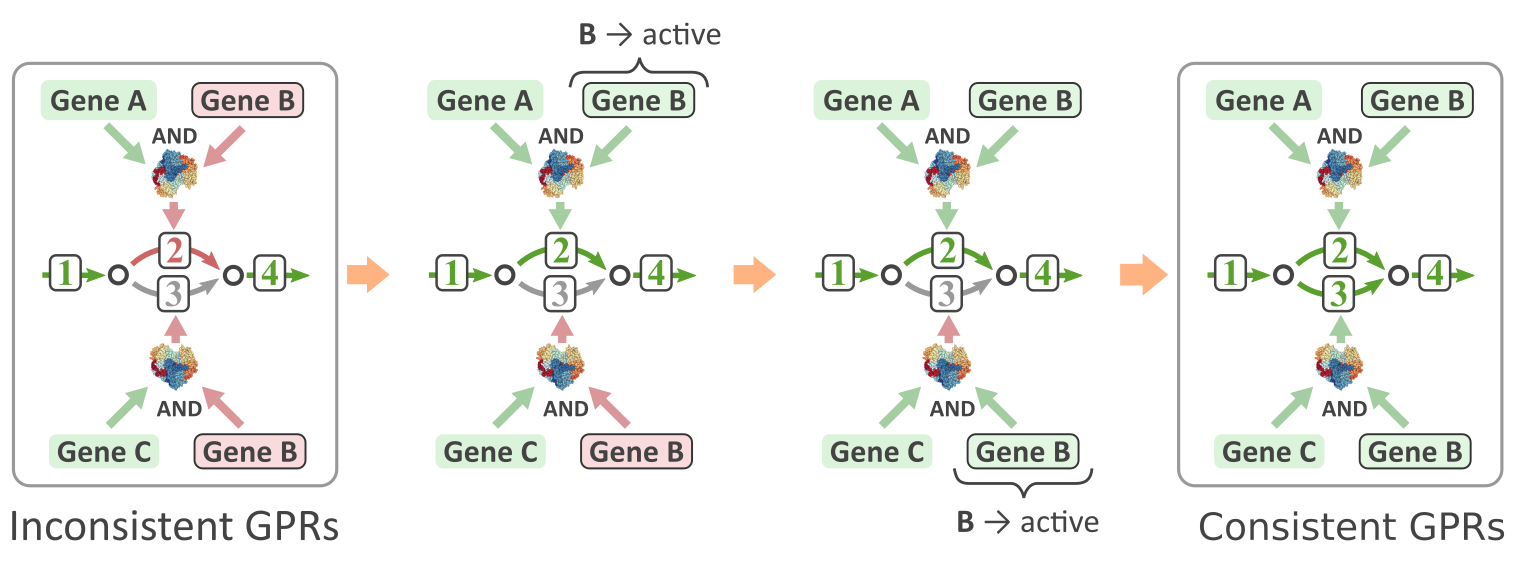

***Figure S3.*** *Toy example consisting of four reactions. Two reactions {1, 4} are categorized as active and the other two {2, 3} as inactive. The CS-Model includes the active reactions and the reaction 2 for gap filling. The inclusion of the gap-filling reaction 2 implies modifying the state of gene B to active. This update must be propagated through the CS-model. Propagating the change of the state of gene B implies that reaction 3 becomes active and thus it should be included in the consolidated CS-Model.*

#### Context-Specific gene knockout

In order to carry out the gene essentially analysis, we performed individual knockouts for each of the genes included in the “corrected reconstructions” (ON state). Note that all the genes not included are treated as inactive (OFF state), since they are not part of the reconstruction.

In order to assess the effect of a gene knockout considering the context we propose the following approach:

1. The confidence score of the knocked out gene is set to its minimum value.
2. Obtain the subset of reactions *A* in which the considered gene is involved.
3. For each reaction in *A*, the confidence score of its associated genes are plugged into the corresponding GPR rules.
4. The GPR rule is solved as previously explained (see *Building a Context-Specific Model* section) to obtain a new confidence score for each reaction in *A.*
5. If the confidence of a particular reaction in *A* becomes OFF, its flux is constrained to 0. Otherwise, its flux bounds are not altered.

After these steps, Flux Balance Analysis (FBA, Orth *et al*, 2010) is used to find the maximum biomass flux in the perturbed system. If the maximum biomass flux predicted drops below a given threshold (in our case less than 1% respect to the wild type value), the deleted gene is predicted to be essential. Otherwise, the gene is predicted to be non-essential.
